# Generation of “nanometer-size aggregates” using Solubility Controlling Peptide tags and their ability to increase a protein’s immunogenicity *in vivo*

**DOI:** 10.1101/793687

**Authors:** Nafsoon Rahman, Mohammad Monirul Islam, Satoru Unzai, Shiho Miura, Yutaka Kuroda

## Abstract

Sub-visible aggregates of proteins are suspected to cause adverse immune response, and a recent FDA guideline has recommended the monitoring of micrometer-size aggregates (2-10 μm) though recognizing that the underlying mechanism behind aggregation and immunogenicity remains unclear. Here, we report a correlation between the immunogenicity and the size of nanometer-scale aggregates of a small 6.5 kDa model protein, Bovine Pancreatic Trypsin Inhibitor (BPTI) variant. BPTI-19A, a monomeric and non-immunogenic protein, was oligomerized into sub-visible aggregates with hydrodynamic radii (*R*_h_) of 3~4 nm by attaching hydrophobic solubility controlling peptide (SCP) tags to its C-terminus. The results showed that the association of non-immunogenic BPTI into nanometer-size aggregates made it highly immunogenic, as assessed by the IgG antibody titers of the mice’s sera. Overall, the study emphasizes that sub-visible aggregates, as small as a few nanometers, which are presently ignored, are worth monitoring for deciphering the origin of undesired immunogenicity of therapeutic proteins.

## Introduction

Therapeutic proteins such as monoclonal antibodies, hormones, and other recombinant protein drugs are increasingly used for the treatment of various types of human diseases. However, protein aggregation, causing unwanted immunogenicity against therapeutic proteins ^**1,2**^ and in turn decreasing therapeutic efficacy through the production of anti-drug antibodies, has occasionally been reported ^**3–5**^. In general, monomeric proteins in any state (native, partially unfolded, or unfolded) may form aggregates of diverse sizes ranging from tiny sub-visible aggregates to visible precipitates with various biophysical properties ^**6–9**^. Hence, it is worth exploring the nature of the relationship between the aggregates and the protein’s immunogenicity from a biophysical perspective.

A link between the aggregation of therapeutic proteins and immunogenicity has long been speculated and has been documented especially for micron-size sub-visible aggregates ^**10–13**^ For example, *in vitro* experiments indicated that 2-10 μm sized sub-visible aggregates of human monoclonal antibodies increased the production of naïve peripheral blood mononuclear cells ^**14**^. Likewise, Trastuzumab (a monoclonal antibody) with less than 3% of ~2 μm sub-visible aggregates, caused T-cell proliferation *in vitro* ^**15**^. *In vivo* investigations are less common, and experiments using mice models showed that sub-visible aggregates of a humanized ScFv generated immunogenicity ^**16,17**^. However, other immunization studies revealed that aggregates of antibodies ^**18**^ or proteins ^**19**^ are not always immunogenic. One factor behind the lack of consistency is that the micron-aggregates are induced by chemical and physical stresses ^**20,21**^ making it difficult to control their biophysical characteristics. Furthermore, the aggregates assessed in samples are different from the injected ones ^**21,22**^ or in many other reports the sizes are not measured at all ^**23–25**^. As a consequence, the biophysical properties (structure, stability, conformation, etc.) of aggregates responsible for the increased immune response are yet to be fully elucidated ^**1,13,17,21**^.

Recently, the European Medicines Agency (EMA) considered visible and sub-visible aggregates as risk factors for unwanted immunogenicity of therapeutic proteins in their safety guidelines ^**26**^. Likewise, the Food and Drug Administration (FDA) recommended the screening and minimizing of sub-visible aggregates within 2-10 μm ranges ^**27**^. These recommendations are based on several clinical reports suspecting that aggregates of therapeutic proteins might have caused adverse immune response. However, clinical case reports are inherently contextual and often put little emphasis on the biophysical properties of the therapeutic proteins ^**28–30**^. Additionally, the immune response generated by protein aggregates smaller than 2 μm is practically un-investigated, due to technical burdens for identifying and measuring them.

In previous reports, we showed, using BPTI-19A as a model protein (58 residues; MW: 5.98 kDa), that one can manipulate a protein’s solubility ^**31–33**^, aggregation kinetics ^**34**^ and aggregate sizes ^**35**^ by using a solubility controlling peptide tag (SCP-tag). The advantage of the SCP-tags is that we can control the size and properties of the aggregates rather accurately and that the tags do not affect or barely affects the thermodynamic properties, structure, and function of the native protein ^**32,33,36**^. In this study, we investigated the immunogenicity of BPTI-19A that were oligomerized into nanometer-sized sub-visible aggregates by attaching hydrophobic SCP-tags (C5A, C5V, C5L and C5I). As a result, the untagged BPTI-19A was not immunogenic as assessed by IgG ELISA against BPTI-19A, but a minute increase of the hydrodynamic radius to ~4 nm, which was generated by attaching a C5I tag to BPTI-19A, increased its immunogenicity by up to 66 folds. These observations indicate that nanometer-size aggregates, which are much smaller than currently examined aggregates and that are difficult to remove by filtration/centrifugation as used in standard biomedical practices, can significantly increase the immunogenicity of a therapeutic protein.

## Results and Discussion

### Oligomerization of BPTI-19A using SCP-tags

The size of the aggregates was measured by dynamic light scattering at both 25 and 37 °C. The samples were prepared in PBS, kept for 20 minutes at 25 °C, and centrifuged at 20,000xg for 20 minutes at 25 °C to remove large particles if any (see Materials and Methods for details). The concentrations of the BPTI variants remained nearly unchanged upon centrifugation or filtration with a 0.2 μm Minisart filter (Supplementary Figure S1a), confirming that only sub-micron and soluble aggregates were present in the samples. The *R*_h_ of the untagged BPTI-19A, C5A, and C5V tagged variants at 25 °C were 1.33±0.02 nm, 1.46±0.05 nm, and 1.34±0.02 nm, respectively, and remained almost unchanged at 37 °C (**Table 1**); the C5L tag increased the *R*_h_ to 2.11±0.14 nm, and the C5I tag increased the *R*_h_ to 3.12±0.06 nm at 25 °C (**Figure 2a** and **Table 1**). All three 5 Ile-tagged variants (N5I, C5I, and ssC5I) formed aggregates with similar hydrodynamic radii (*R*_h_ ~3.12 nm to ~3.97 nm) at 37 °C (**Table 1** and Supplementary Figure S1b). Importantly, the sizes of the sub-visible aggregates analyzed just before immunization remained almost constant from dose to dose (**Figure 2b**). These observations clearly suggested that the increased hydrodynamic radii observed for the C5I-tagged BPTI originated from the SCP-tag attachment. The overall order of hydrodynamic radii at 37 °C was thus BPTI-19A<C5A/C5V<C5L<N5I<C5I/ssC5I (**Table 1** and Supplementary Figure S1b) fully corroborating our previous observations ^**35**^. Similarly, static light scattering (SLS) showed that the scattering intensities increased with increasing hydrophobicity ^**37**^ of the SCP-tags (**Figure 2c** and Supplementary Figure S1c) indicating that the attachment of a hydrophobic Ile-tag produced aggregates which are in agreement with DLS observations (Supplementary Figure S1d).

**Table 1:** Hydrodynamic radius of BPTI-19A and its SCP-tagged variants. Proteins were formulated at 0.3 mg/mL concentrations in PBS. DLS measurements were carried out just before immunization at 25 °C followed by 37 °C and again at 25 °C (re-25 °C). The values are the average of three independent measurements where ± sign indicates a standard deviation (SD). *R*_h_ was calculated from the number distributions using the Stokes-Einstein equation.

| Mutant Identities | Hydrodynamic Radius ($R_h$, nm) in PBS | | |
| --- | --- | --- | --- |
|  | 25 °C | 37 °C | re-25 °C |
| BPTI-19A | 1.33±0.02 | 1.34±0.02 | 1.26±0.016 |
| BPTI-C5A | 1.46±0.05 | 1.53±0.06 | 1.43±0.06 |
| BPTI-C5V | 1.34±0.02 | 1.39±0.06 | 1.31±0.06 |
| BPTI-C5L | 2.11±0.14 | 2.34±0.13 | 2.19±0.26 |
| BPTI-N5I | 1.49±0.30 | 3.13±0.23 | 2.85±0.44 |
| BPTI-C5I | 3.12±0.06 | 3.71±0.13 | 3.23±0.10 |
| BPTI-ssC5I | 3.47±0.09 | 3.97±0.22 | 3.42±0.08 |

**Figure 1.**
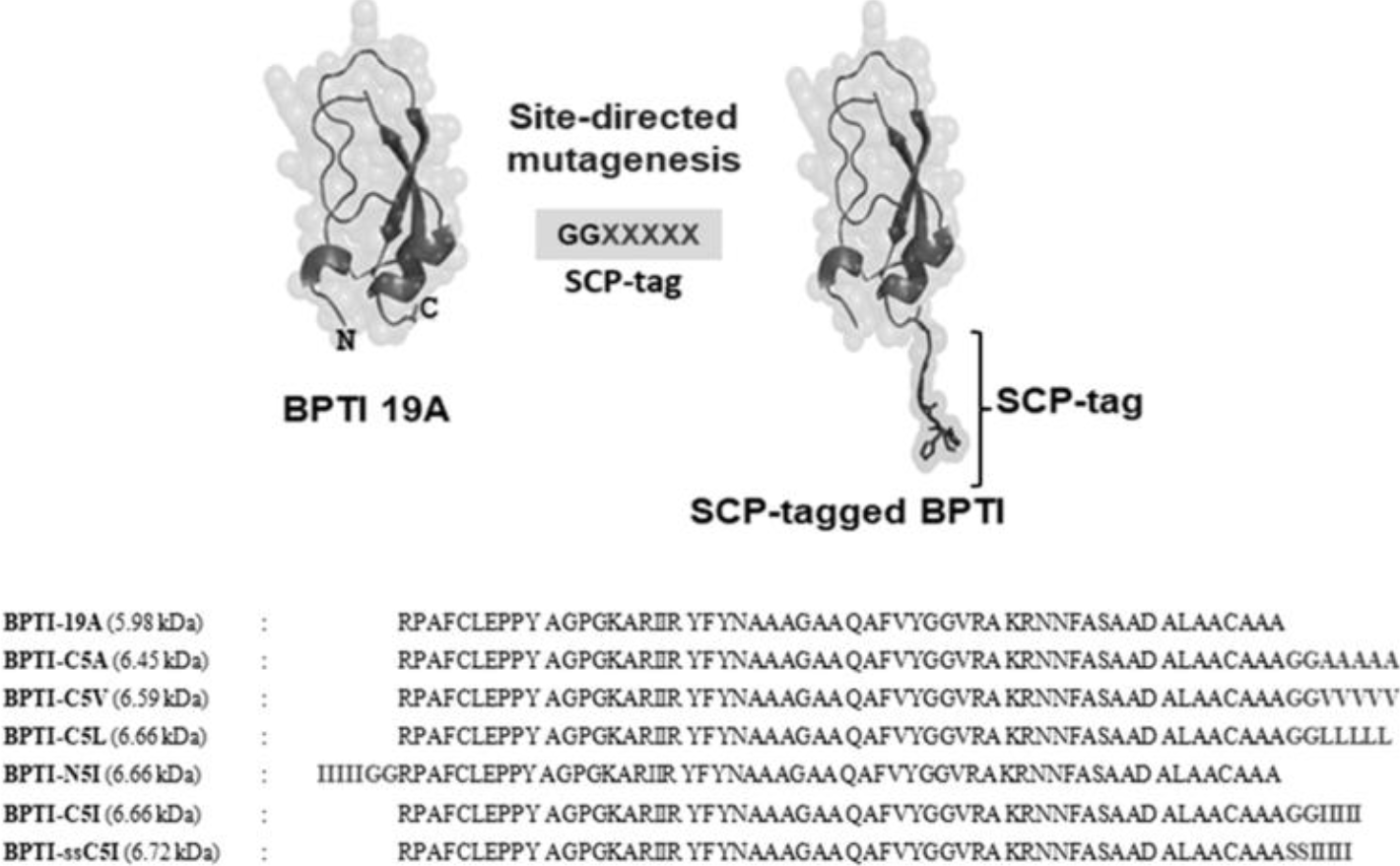
Sequences and structure of SCP-tagged BPTI. The untagged BPTI-19A contains 58 amino acid residues. C5A, C5V, C5L, and C5I variants were designed by attaching their respective tags at the C-terminus (site-directed mutagenesis) except N5I having the tag at the N-terminus. Two glycine or serine residues served as a spacer between the host protein and the SCP-tags.

**Figure 2.**
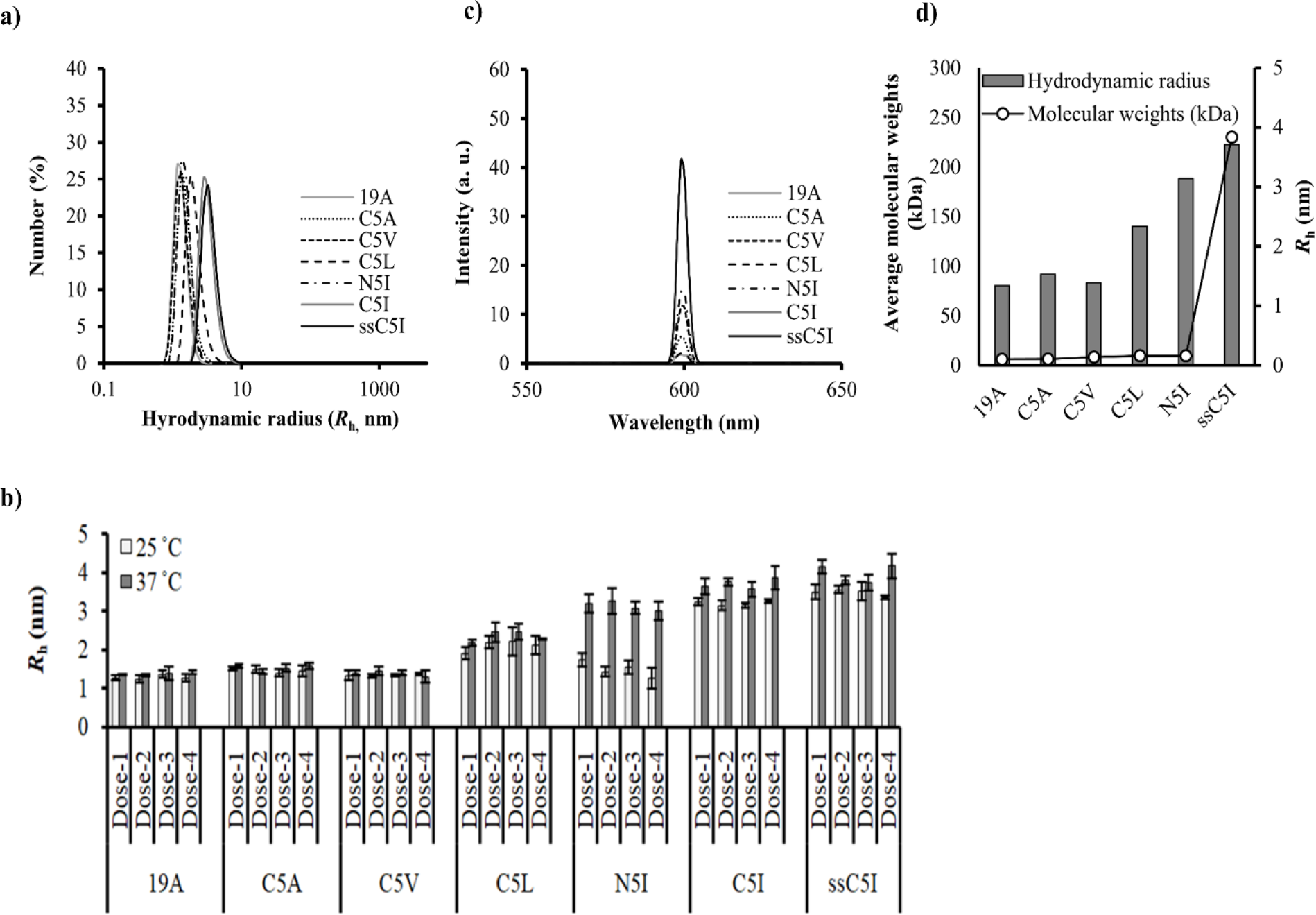
Influence of SCP-tags on sub-visible aggregates’ sizes measured by DLS and SLS and AUC. a) DLS spectra of the size distribution Number (%) at 25 °C. b) Hydrodynamic radii of SCP-tagged BPTI variants measured just before immunization from dose-1 to dose-4 at 25 (white bars) and 37 °C (grey bars) respectively. The *R*_h_ were computed from DLS’s number spectra. c) Aggregation intensities of SCP-tagged BPTI variants at 25 °C measured by SLS. d) Increase in the average molecular weights at 33 °C (black line) with respect to the hydrodynamic radii (bars) of SCP-tagged BPTI variants at 37 °C as measured by AUC. Proteins were formulated at 0.3 mg/mL concentrations in PBS pH 7.4 for all measurements. Values are shown as the average of three independent measurements and three accumulations for DLS and SLS, respectively. Line symbols are explained within the panels.

In addition to DLS and SLS, the sizes of the aggregates were determined by sedimentation velocity experiment using analytical ultra centrifugation (AUC). The results showed that at 33 °C BPTI-19A and C5A had homogenous average molecular weights of ~6 kDa indicating that they are monomers. C5V was also nearly homogenous with an average molecular weight of ~8 kDa (**Figure 2d** and **Table 2**). However, C5L and N5I contained an oligomeric populations. In particular, ssC5I, which had the largest *R*_h_ showed various species of aggregates existed in the solution, and the average molecular weight of the aggregates was estimated to be ~230 kDa. Altogether, AUC and the SLS results were in good agreement with the DLS observations where C5I tag induced the highest oligomeric size, intensity and molecular weight.

**Table 2:** Sedimentation velocity experiment with BPTI-19A and its SCP-tagged variants using AUC. AUC experiments were carried out with a protein concentration of 0.3 mg/mL at 33 °C. The c (s) distribution was converted into a molar mass distribution c (M) and thus the average molecular weights of different BPTI variants were calculated. C5I could not be measured as it contained various populations (indicated by ‘*’).

| **Mutant** | **Molecular weights (kDa)** | **Population (%)** |
| --- | --- | --- |
| **Identities** | **33 °C** | **33 °C** |
| BPTI-19A | 6.1 kDa | 89.66% |
| BPTI-C5A | 6.3 kDa | 92.13% |
| BPTI-C5V | 8.3 kDa | 74.59% |
| BPTI-C5L | 9.7 kDa | 48.37% |
| BPTI-N5I | 9.6 kDa | 56.98% |
| BPTI-C5I | Large* | * |

### Effects of SCP-tags on the structures of BPTI variants

The impacts of SCP-tags on the conformational stability of BPTI-19A were investigated using Tyr-fluorescence and CD under conditions identical to those used for DLS (0.3 mg/mL in PBS, pH 7.4). Fluorescence spectra showed that all BPTI variants maintained native conformation except C5I/ssC5I which quenched the tyrosine fluorescence at 25 and 37 °C (**Figure 3a** and **3b**, Supplementary Figure S2a). Additionally, the fluorescence of C5L showed a marginal reduction when the temperature was increased to 37 °C. We can interpret this quenching as a partial denaturation, where the tyrosine residues are becoming exposed to water molecules and their mobility increase ^**38**^ Note that, all the variants showed reversibility when the temperature was cooled back to 4 °C (Supplementary Figure S2b–S2c) which corroborated with our previous results ^**35**^.

**Figure 3.**
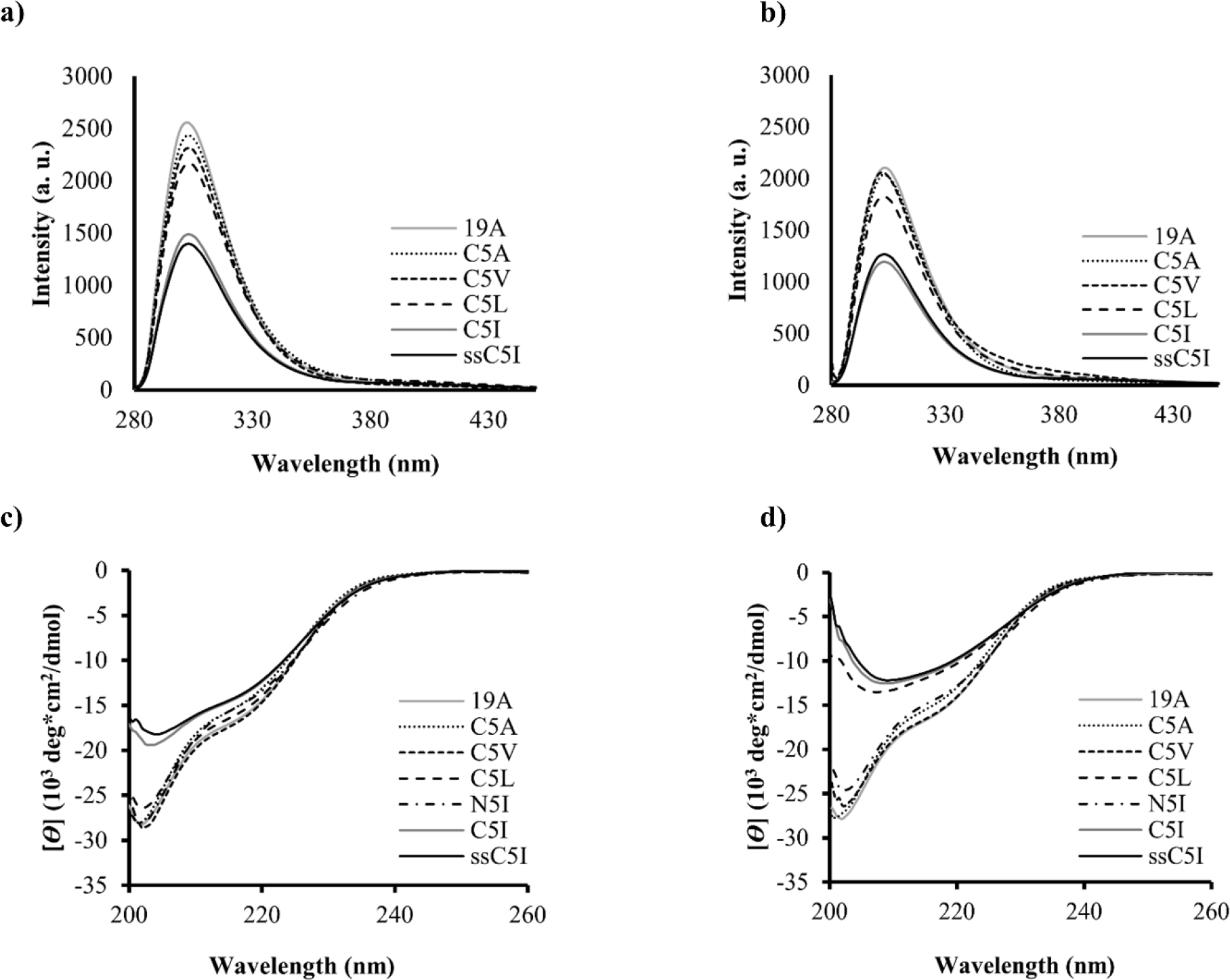
Spectroscopic analysis of untagged BPTI-19A and its SCP-tagged variants by fluorescence and CD measurements. Tyr-fluorescence spectra of 19A, C5A, C5V, C5L and C5I/ssC5I at a) 25 °C and b) 37 °C. CD spectra of all BPTI variants at c) 25 °C and d) 37 °C. All protein samples were formulated in PBS, pH 7.4, at a concentration of 0.3 mg/mL. Three accumulations were taken for each measurement. Line symbols are explained within the panels.

To understand the effect of secondary structural changes on the conformation of BPTI variants CD measurements were carried out and the supernatants’ spectra at 25 and 37 °C revealed that none of the SCP-tags affected the native-like secondary structure contents, except C5I/ssC5I (**Figure 3c** and **3d,** Supplementary Figure S2d-S2j) which was slightly denatured at 25 and 37 °C (C5L was partially denatured at 37 °C but not at 25 °C in line with the above tyrosine fluorescence results). This partial denaturation occurs because of the hydrophobic environment created by the hydrophobic tags upon BPTI-19A’s oligomerization and we related this to a multi-molecular reversed hydrophobic effect ^**39**^. The association was reversible when CD spectra were measured at low concentration or temperature, and the tagged BPTI variants are fully folded as previously assessed by Trypsin inhibition assay performed at 280 nM ^**35**^. Thus, the present results confirm the reversible association of all SCP-tagged BPTI-19A variants, which retain a native-like structure in the monomeric state, and they suggest that the SCP-tags did not affect the native biophysical properties of BPTI-19A, except producing temperature-dependent nanometer-size sub-visible aggregates under inoculation condition used in the present study.

### Effects of nanometer size aggregates on protein immunogenicity

We investigated the sizes of SCP-tag induced sub-visible aggregates on the immunogenicity of BPTI-19A in Jcl: ICR mice by ELISA. Both in the presence and absence of adjuvant, the untagged BPTI-19A with *R*_h_ ~1.34 nm was essentially non-immunogenic (Supplementary Figure S3a), as expected for a small protein. Similarly, no immune response was observed during the first two doses for any of the variants even in the presence of adjuvant. However, after the third dose, the antibody titers of the tagged BPTIs started to increase (**Figure 4a**), and after the final dose, the heart-bleed sera of C5I/ssC5I exhibited the highest absorbance at 492 nm (**Figure 4b**). C5A and C5V having *R*_h_ close to untagged 19A (*R*_h_ ~1.5 nm) and C5L which was slightly larger (*R*_h_ ~2.3 nm) than that of the untagged BPTI-19A, increased the immunogenicity by 15-23 fold. Upon a further increase of the hydrodynamic radius to ~3.7 nm by attaching the 5-Ile tags, the anti-BPTI-19A titers increased over 55 fold (**Table 3** and Supplementary Figure S3b), in line with their aggregate sizes and SLS intensities (**Figure 4c** and Supplementary Figure S3c). Notably, ssC5I (*R*_h_ ~3.96 nm) also enhanced the titer by 66 fold indicating that the increase in antibody titers observed with 5-residue Ile-tag was independent of the type of residues (Gly or Ser) used as a spacer. On the other hand, the N-terminus-tagged BPTI-N5I increased the IgG titer only 22 fold which was almost one-third of the average titer observed for C5I (N5I also formed smaller aggregates than C5I, *R*_h_ ~3.13 nm). Noteworthy, the antibody responses were directed against the BPTI-19A and not against the tag sequence as demonstrated by the ELISA signals, obtained with plates coated with BPTI-19A without a tag or BPTI-19A with their respective tags, self-tags. (Supplementary Figure S3d). Immunization experiments in the absence of adjuvant showed similar results, in which C5I increased the antibody titer of BPTI-19A by 90 to 222 fold (**Figure 4d** and **Table 3**). Thus, the effect of the tags was comparable, and furthermore cumulative, to that of the traditional Freund’s adjuvant (Supplementary Figure S3e), confirming the strong potential of SCP-tag as a target-specific adjuvant.

**Table 3:** Average IgG titer values and fold-increase of antibody titers for SCP-tagged BPTI variants with and without adjuvant. ^1^ The titers were calculated using a power fitting model, and the values were averaged using the number of the mice (n) in respective groups [19A (+), n=4; C5A/V/L (+), n=3; N5I (+), n=4; C5I (+) n=3; ssC5I (+), n=4; 19A (−), n=3; C5I (−), n=3; ssC5I (−), n=3]. ^2^ Fold-increase of antibody titers in the presence and absence of adjuvant were calculated with respect to the titer of BPTI-19A (+) and BPTI-19A (−), respectively. ‘×’ indicates the absence of tags.

| Dose formulation | Mutants | Tags | Average IgG Titers<sup>1</sup> | Fold increased<sup>2</sup> |
| --- | --- | --- | --- | --- |
| With adjuvant (+) | BPTI-19A | × | 1171.9 | 1 |
|  | BPTI-C5A | Gly<sub>2</sub> Ala<sub>5</sub> | 27280.3 | 23.3 |
|  | BPTI-C5V | Gly<sub>2</sub> Val<sub>5</sub> | 18251.5 | 15.6 |
|  | BPTI-C5L | Gly<sub>2</sub> Leu<sub>5</sub> | 18467.1 | 15.8 |
|  | BPTI-N5I | Ile<sub>5</sub> Gly<sub>2</sub> | 26071.8 | 22.2 |
|  | BPTI-C5I | Gly<sub>2</sub> Ile<sub>5</sub> | 64846.8 | 55.3 |
|  | BPTI-ssC5I | Ser<sub>2</sub> Ile<sub>5</sub> | 77825.5 | 66.4 |
| Without adjuvant (-) | BPTI-19A | × | 82.1 | 1 |
|  | BPTI-C5I | Gly<sub>2</sub> Ile<sub>5</sub> | 18267.1 | 222.5 |
|  | BPTI-ssC5I | Ser<sub>2</sub> Ile<sub>5</sub> | 7417.4 | 90.4 |

**Figure 4.**
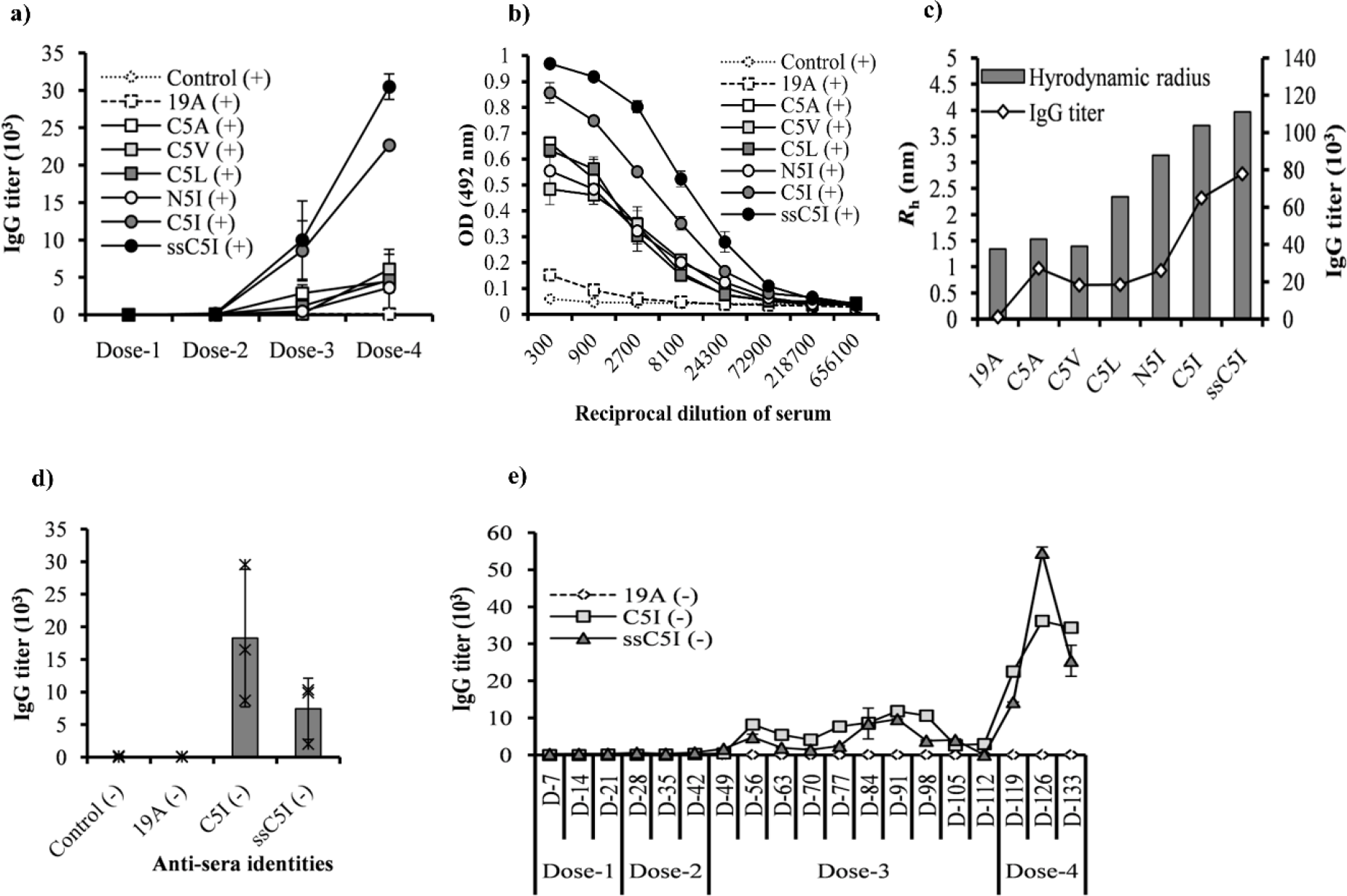
Antibody responses against untagged BPTI-19A and its SCP-tag induced sub-visible aggregates assayed by ELISA. Immunization in the presence of adjuvant is indicated by ‘+’, whereas ‘-’ stands for immunization in the absence of adjuvant. a) Dosewise antibody titers of untagged BPTI-19A and SCP-tagged BPTI variants (C5A, C5V, C5L, N5I, and C5I/ssC5I) in the presence of adjuvant using tail-bleed serum samples. b) OD at 492 nm of anti-BPTI-19A, C5A, C5V, C5L, N5I, and C5I/ssC5I sera (with adjuvant) measured by ELISA of the respective heart bleed samples. c) Increase in antibody titers (black line) with respect to the hydrodynamic radii (bars) of SCP-tagged BPTI variants at 37 °C. d) Antibody titers produced against untagged-19A and C5I/ssC5I-tagged BPTIs in the absence of adjuvant. ‘*’ shows the results for individual mice. e) Long term antibody titers against BPTI-19A and BPTI-C5I/ssC5I in the absence of adjuvant. ‘D’ indicates the number of days at which tail bleeding was performed counting from the first inoculation. All doses were formulated at 0.3 mg/mL concentrations in PBS throughout the immunization scheme. Line symbols are explained within the panels.

### Immune response maintenance and immunogenic memory

Finally, we assessed the long-term maintenance of the immune responses generated by the untagged BPTI-19A and BPTI-C5I. Mice were injected with three consecutive doses at 3-week intervals, kept untreated for 70 days (10 weeks), but during this period the IgG level was measured weekly through tail-bleeding. Anti-BPTI-19A IgG levels, as measured by ELISA, remained high over 8 weeks for the mice injected with C5I, before declining. Furthermore, BPTI-19A memory was generated by the aggregates. Seventy days after the 3^rd^ dose, the 4^th^ dose of C5I boosted the IgG titers by 2300-3600 fold, whereas the response of mice immunized with the untagged BPTI-19A remained almost nil throughout the entire immunization scheme (**Figure 4e**). Altogether, these observations indicated that the immune response was long-lasting and further that B-cell immune memory was established.

### SCP-tags for biophysical and immunological study

A link between protein aggregation and immunogenicity has been suspected for a long time, however, without reaching a strong consensus. The lack of consistency was often blamed on to the inherent disparity arising from *in vivo* experiments, but it appears that in many reports, the sample condition might not have been sufficiently controlled or even measured. From this viewpoint, the nanometer-size sub-visible aggregates produced by attaching hydrophobic SCP-tags, are well controlled and avoid the random events encountered by aggregates produced by physical or chemical stresses. Furthermore, we carefully assessed the aggregates’ size and its stability upon temperature, pH, protein concentration and type of tags. We further performed a near “real-time” monitoring of the hydrodynamic radius of the aggregates ^**40**^ just before each round of inoculation. We believe that such monitoring of the aggregates’ size is essential as these aggregates may grow and dissociate upon small variation of external factors. Consequently, SCP-tags combined with our “real-time” monitoring protocol could provide a new tool for producing aggregates in a stable manner and analyzing the effect of aggregation on immunogenicity, from a biophysical viewpoint.

### Conclusion

The effects of sub-visible aggregates of a few nanometers dimension on a protein’s immunogenicity have mostly been ignored due to technical challenges in isolating, characterizing, and monitoring them. To the best of our knowledge, this is the first report showing as directly as currently possible, that aggregates as small as a few nanometers (*R*_h_ <5 nm) can convert a non-immunogenic protein into a highly immunogenic one. Aggregates smaller than 0.2 μm (200 nm) cannot be removed by filtering using standard filters or centrifugation, and our study thus emphasizes the usefulness of monitoring the presence of small oligomers (or sub-visible aggregates) during the storage and formulation of therapeutic proteins. Finally, the present results show that the SCP-tags can be used to control the aggregates’ size in a systematic and stable manner, and as such, they can contribute to a better understanding of the mechanisms underlying aggregation-triggered immunogenicity of proteins.

## Materials and Methods

### Construction of SCP-tagged BPTI variants (C5A, C5V, C5L, C5I/ssC5I and N5I)

All BPTI variants were constructed with a pMMHa vector ^**35**^. DNA sequences encoding two Gly/Ser residues (serving as a spacer), and the SCP-tags were added to the C-terminus of the template BPTI-19A [a BPTI containing 19 alanines out of its 58 residues ^**31**^ sequence using QuikChange site-directed mutagenesis (Stratagene, USA). The plasmid sequences were confirmed by DNA sequencing (ABI PRISM 3130xl Genetic Analyzer, USA). BPTI variants were named according to the number and type of amino acids added to the C-terminus of BPTI-19A **(Figure 1)**. For example, C5I stands for five Isoleucine residues added after two glycines as a spacer at the C-terminus of BPTI-19A. Similarly, N5I indicates a mutant with five isoleucine added at the N-terminus of BPTI with two glycines as a spacer.

### Protein expression, purification and sample preparation

All BPTI variants were expressed in *Escherichia coli* BL21 (DE3) pLysS cell lines at 37 °C and purified by pI precipitation, followed by reverse phase HPLC as per our previous protocol ^**35**^. Protein identities were confirmed by MALDI-TOF mass spectrometry (AB SCIEX TOF/TOF TM 5800, USA) and the purified proteins were preserved at −30 °C as lyophilized powder. While preparing samples, the lyophilized protein powders were first dissolved at 1mg/mL in MilliQ water (Millipore A10 ultra-pure water purifier, EMD Millipore, Germany) as protein stocks, and the final working sample was prepared in phosphate buffered saline (PBS, pH 7.4-filtered by 0.2 μm Minisart filters) at 0.3 mg/mL concentration as determined by the extinction coefficient (A_280_) using Nanodrop (Nanodrop-2000, Thermo Fisher Scientific, USA). Stock samples were centrifuged at 20,000xg for 20 minutes at 4 °C. Working samples were kept at 25 °C for 20 minutes and centrifuged just before measurements to remove aggregates, if there was any.

### Analysis of sub-visible protein aggregates by dynamic light scattering (DLS), static light scattering (SLS) and analytical ultra-centrifugation (AUC)

The sub-visible aggregates’ sizes were measured by preparing the BPTI variants in PBS at 0.3 mg/mL. As mentioned earlier, samples were left still for 20 minutes at 25 °C, centrifuged (20,000xg for 20 minutes at 25 °C), and the supernatants were used for DLS measurements (Malvern Zetasizer Nano-S System, UK). The *R*_h_ was measured at 25 and 37 °C and calculated as the average of three independent readings from the number distributions using the Stokes-Einstein equation ^**41**^. The presence of sub-visible aggregates was also monitored using static light scattering (SLS). The intensity of the supernatants was monitored at 600 nm wavelength at 25 and 37 °C with an FP-8500 spectrofluorometer (JASCO, Japan) using a cuvette with a 3 mm optical path length. Each measurement was accumulated three times, and the values were averaged.

AUC’s samples were prepared in the same way as the injected samples (or DLS) at a protein concentration of 0.3 mg/mL. Sedimentation velocity experiments were carried out using an Optima XL-A analytical ultracentrifuge (Beckman-Coulter, USA) with a four-hole An60Ti rotor at 25 and 33 °C. Before centrifugation, the samples were dialyzed overnight against buffer solution with PBS pH 7.4. The solvent density, viscosity and protein partial specific volumes were calculated using SEDNTERP software ^**42**^. Each sample was then transferred into a cell with a two-channel centerpiece. Data were obtained at 50,000 rpm. Absorbance at 280 nm scans were collected at 10 minutes intervals during sedimentation. Data analysis was performed by continuous distribution *c* (s) analysis module in the SEDFIT program ^**43**^. Frictional ratio (f/fo) was allowed to float during fitting. The *c* (s) distribution was converted into a molar mass distribution c (M).

### Structural analysis of BPTI variants by Tyr Fluorescence and Circular Dichroism (CD) spectroscopy

Tertiary and secondary structures of all BPTI variants were monitored through Tyr-fluorescence and CD spectroscopy using an FP-8500 spectrofluorometer and a JASCO J820 CD spectropolarimeter (JASCO, Japan), respectively. Tyr-fluorescence (λ_ex_ 275 nm) of all BPTI variants was measured at various temperatures such as 4-25-37 °C, and the reversibility was checked by cooling the sample to 4 °C (re- 4 °C). A quartz cuvette with 3 mm optical path length was used to carry out the fluorescence measurements, and *λ*_em_ was recorded between the wavelengths of 280-450 nm. Similarly, the CD spectra of proteins’ supernatants were recorded in the wavelength range of 200-260 nm using a 1mm optical path length cuvette at 25 and 37 °C and cooling them back to 25 °C (re- 25 °C). In both experiments, the concentrations of the protein samples were maintained as same as the DLS or SLS measurement conditions (0.3 mg/mL in PBS pH 7.4.)

### Immunization experiments

The inoculation samples were prepared by dissolving the lyophilized powder in MilliQ water at a concentration of 1 mg/mL. The samples were aliquoted into 10 single-use samples and kept at −30 °C until use. Prior to each inoculation, the samples were slowly thawed at room temperature, reconstituted into the final buffer (PBS) at a concentration of 0.3 mg/ml, incubated for 20 minutes at 25 °C, centrifuged, and the aggregates’ sizes were monitored by measuring the *R*_h_ at 25 and 37 °C by DLS (see above for measurement details).

Four-week-old female mice (Jcl: ICR, CLEA, Japan) were injected with the BPTI variants. All of the experiments were performed in compliance with TUAT’s and Japanese governmental regulations on animal experimentation. The mice were accommodated on sterilized wood bedding at an ambient temperature maintained at 22±0.5 °C and relative humidity around 47±8% with a natural half-day light-dark cycle. Subcutaneous (SC) injection was carried out with the untagged and SCP-tagged BPTIs at 30 μg/100 μL/dose without adjuvant. Additionally, for injection with adjuvant (WAKO, Japan), the first dose was given subcutaneously with Freund’s incomplete adjuvant (protein: adjuvant = 1:1), and doses 2-4 with Freund’s complete adjuvant were given intraperitoneally (IP) at weekly intervals. In all experiments, the quantity of injected protein was 30 μg. Control mice were injected with PBS in the presence or absence of adjuvant.

Three days after each inoculation, ~20 μL of tail-bleed samples were collected and used for measuring the level of anti-BPTI IgG antibodies by using ELISA (see the next section). After the final dose, heart-bleed samples were collected when mice were sacrificed through spinal-cord dislocation. These heart-bleed serum samples were preserved in 1:4 dilutions in PBS supplemented with 20% glycerol and stored at −30 °C.

### Detection of anti-BPTI responses by ELISA

Anti-BPTI immune response was measured by ELISA using 96-well microtiter plates (TPP micro-titer plates, Japan). HPLC purified untagged BPTI-19A (2.5 μg/mL in PBS) was used for overnight coating of the plate at 25 °C. Unbound proteins were washed out with PBS, and the plates were blocked with 1% BSA in PBS for 45 minutes at 37 °C, followed by washing with PBS once.

Anti-BPTI sera were applied at an initial dilution of 1:50 for the tail-bleed samples, and 1:300 for the heart-bleed samples, followed by 3-fold serial dilution in subsequent wells containing 0.1% BSA-PBS. Plates were then incubated at 37 °C for 2 hours. A positive control was used in all ELISA plates. After washing three times with PBS-0.05% Tween-20, 100μl/well of anti-mouse IgG HRP conjugate (Thermo Fisher Scientific, USA; at 1:3000 dilution in 0.1% BSA-PBS-Tween-20) was added and incubated at 37 °C for 90 minutes. Finally, unbound conjugates were removed by thoroughly washing three times with PBS-0.05% Tween-20. Coloring was performed by adding the O-phenyl Di-amine (OPD) substrate and incubating for 20 minutes (100 μl/well; OPD 0.4 mg/ml supplemented with 4 mM hydrogen peroxide). Color intensities were measured at 492 nm using a microplate reader (SH-9000 Lab, Hitachi High-Tech Science, Japan) immediately after stopping the reaction with 1 N sulfuric acid (50 μL/well).

## Acknowledgements

We thank the members of the Kuroda’s laboratory for technical help and discussion. We are grateful to Ms. Patricia S. McGahan for English proofreading and Profs Tsuyoshi Tanaka and Tomoko Yoshino and for the use of the Zetasizer.

## Author Contribution

Y.K., M.M.I., N.R. designed the project, and wrote the manuscript. N.R., S.U., M.M.I., and S.M. performed the experiments and analyzed and compiled the data. All authors read and approved the manuscript.

## Data Sharing

All data are given in the manuscript and the supplementary data.

## Notes

The SCP-tag sequences are covered by a Japanese patent 5273438 and an international (PCT) patent application PCT/JP2018/029395.

## Supplementary Information

**Figure S1:**
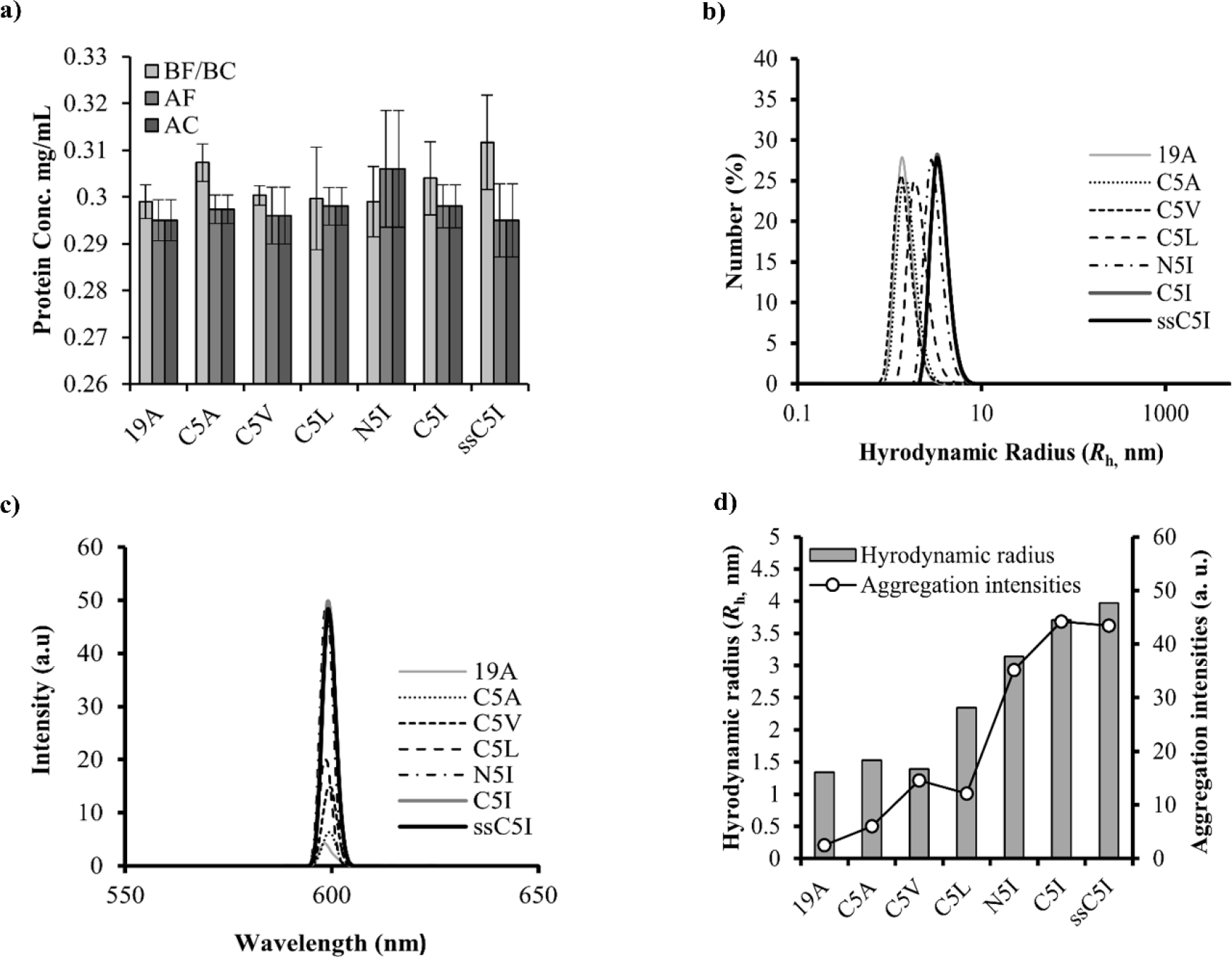
**a)** Effect of filtration and centrifugation on the concentrations of SCP tagged variants in at 0.3 mg/ml concentration PBS pH 7.4. BF/CF means the concentration ‘before filtration or centrifugation’ and AF/AC means ‘after filtration or centrifugation’. Three measurements were carried out and the values were averaged **b)** Hydrodynamic radius of untagged BPTI-19A and SCP tagged variants (0.3mg/mL in PBS pH 7.4) at 37 °C in terms of their number (%). Line symbols are explained within the panels **c)** Aggregation intensities of SCP tagged variants (0.3mg/mL in PBS pH 7.4) at 37 °C measured by SLS. Three accumulations were carried out for each mutant. Line symbols are explained within the panels **d)** Increase in aggregation intensities of SCP tagged BPTI variants measured by SLS (0.3 mg/mL in PBS pH 7.4) at 37 °C with respect to their hydrodynamic radii.

**Figure S2:**
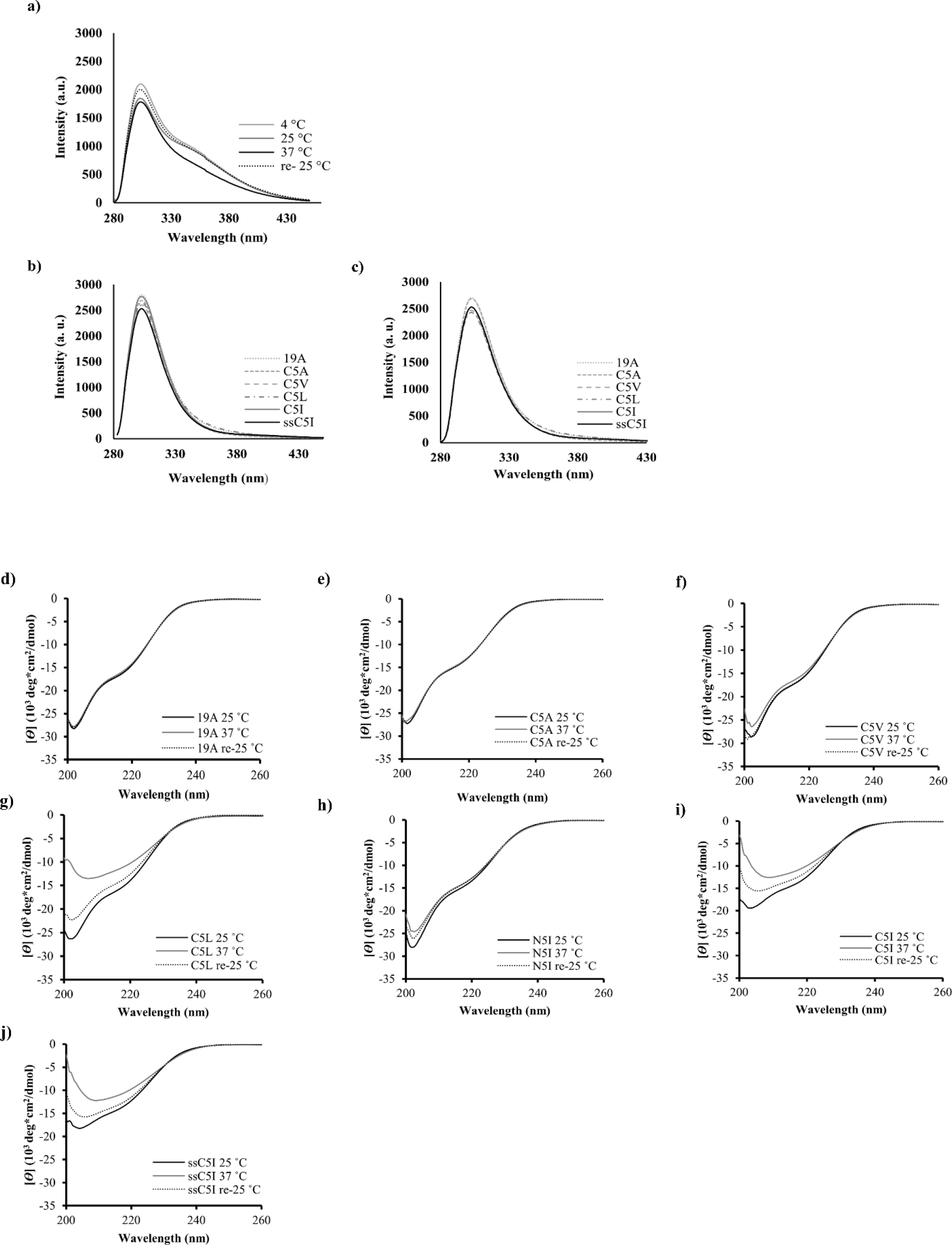
Tyr-fluorescence of **a)** N5I at 4-25-37 and re- 4 °C (after cooling back from 37 °C) **b)** Fluorescence spectra of 19A, C5A, C5V, C5L, C5I and ssC5I at **b)** 4 °C and **c)** re- 4 °C (after cooling back from 37 °C. CD analysis of the secondary structure contents of individual BPTI variants (**d)-(j)** was measured at 25, 37 and re- 25°C (after cooling back from 37 °C). Three accumulations were carried out for each mutant for both fluorescence and CD measurements with a protein concentration of 0.3 mg/mL in PBS pH 7.4.

**Figure S3:**
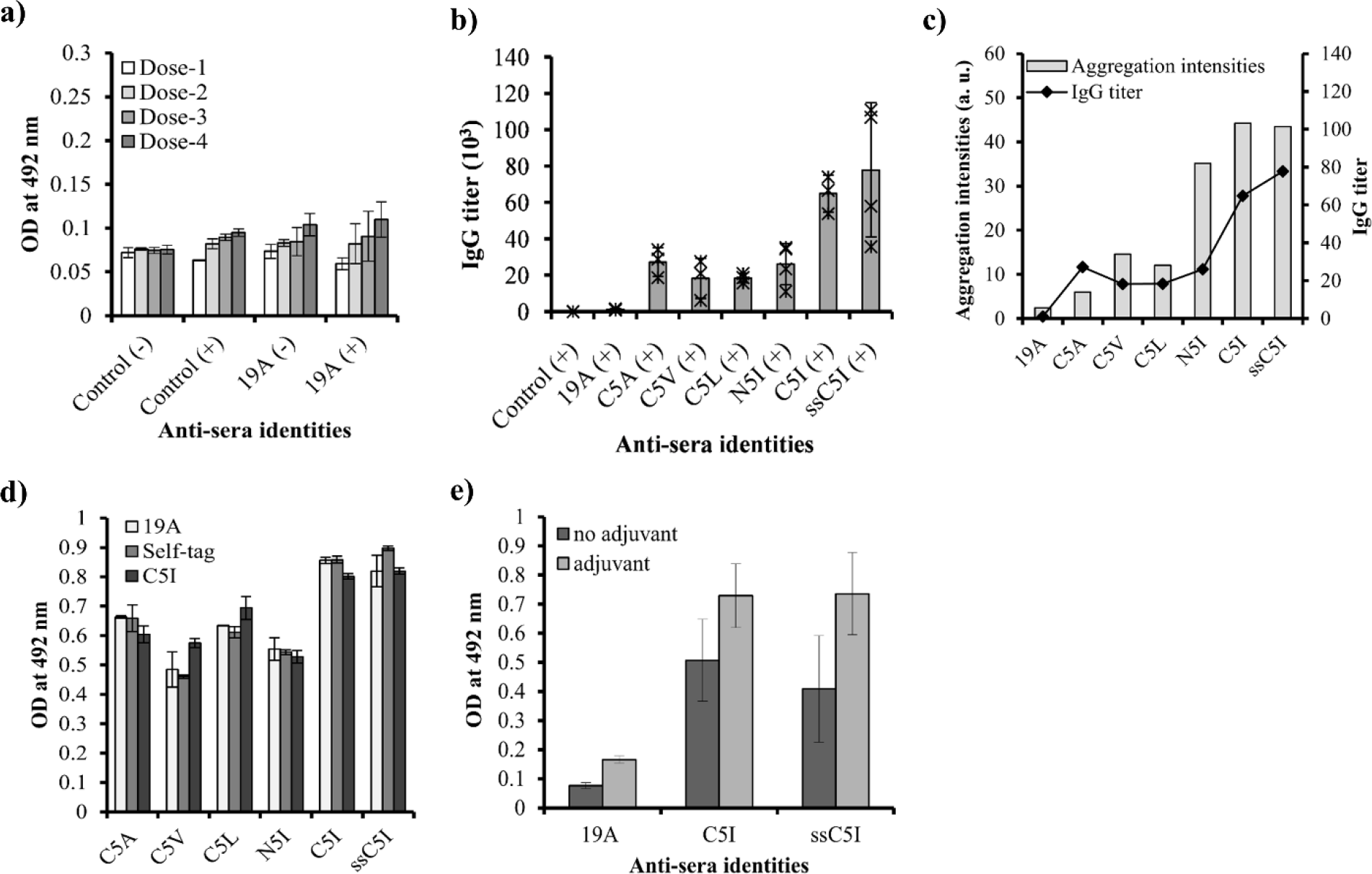
**a)** Dose dependent OD values of control-sera (raised against PBS) and untagged BPTI-19A at 492 nm in the presence (+) and absence (−) of adjuvant **b)** Antibody titers of BPTI-tagged variants in the presence of adjuvant (+) after sacrificing the mice. Number of mice were indicated by ‘*’ on the bar diagrams **c)** Increase in antibody titers of SCP tagged BPTI variants with respect to their aggregation intensities measured by SLS at 37 °C **d)** OD values of sera raised against C5A, C5V, C5L, N5I and C5I/ssC5I against coating antigen untagged BPTI-19A, self-tags and C5I **e)** OD values of heart-bleed sera of untagged BPTI-19A and BPTI-C5I/ssC5I at 492 nm in the presence (+) and absence (−) of adjuvant.

